# Walk This Way: Modeling Foraging Ant Dynamics in Multiple Food Source Environments

**DOI:** 10.1101/2024.01.20.576461

**Authors:** Sean Hartman, Shawn D. Ryan, Bhargav R. Karamched

## Abstract

Foraging for resources is an essential process for the daily life of an ant colony. What makes this process so fascinating is the self-organization of ants into trails using chemical pheromone in the absence of direct communication. Here we present a stochastic lattice model that captures essential features of foraging ant dynamics inspired by recent agent-based models while forgoing more detailed interactions that may not be essential to trail formation. Nevertheless, our model’s results coincide with those presented in more sophisticated theoretical models and experiment. Furthermore, it captures the phenomenon of multiple trail formation in environments with multiple food sources. This latter phenomenon is not described well by other more detailed models. An additional feature of this approach is the ability to derive a corresponding macroscopic PDE from the stochastic lattice model which can be described via first principle interactions and is amenable to analysis. Linear stability analysis of this PDE reveals the key biophysical parameters that give rise to trail formation. We also highlight universal features of the modeling framework that this simple formation may allow it to be used to study complex systems beyond ants.

## 1. Introduction

Social insect behavior has fascinated the biological community for a long time. The collective behavior of insects can be used to achieve more complicated outcomes than possible for individuals. This is especially important for ant species that rely on communication via pheromone to coordinate activity. In the absence of an external motivation or central control center, ants can perform complex sets of tasks [30, 31] and exhibit macroscopic emergent behavior [24, 27]. This makes them an ideal model organism for studying the physical origins of self-organizing behavior. For example, in order to find food for survival, ant colonies send foragers away from the nest executing a random search process [57]. Once food is found the ants secrete a pheromone to attract other nearby foragers which eventually leads to the formation of a trail (e.g., [24, 32, 36, 38, 39, 52, 57, 68] among others). The ants proceed to break down the food source until interruption (i.e., by a predator or weather) or completion before returning to the colony.

The trail formation is a rich source of collective behavior observations as ants self-organize into lanes along the trail for optimal resource transport. While fascinating to watch, there is still much to be understood about the raiding process—specifically, the raiding process in the presence of multiple food sources. What makes this challenging to model is that individuals within an ant colony cannot directly communicate and instead rely on chemical signaling through pheromones.

Thus, in the presence of multiple signals how will the colony proceed through the raiding process to ensure an efficient acquisition of resources?

A given colony of ants provides an ideal biological system to study self-organization because all participating agents are identical and have the same motivations: discover food and break it down for retrieval [12]. The results derived here provide foundational insight into similar social insect systems such as locusts [5, 33] and termites [18]. Though detailed observations of ant foraging behavior have become increasingly available over the last century (cf [60]), mathematical models are still striving to uncover the underlying behavior of this natural phenomenon. The desire to develop a mathematical model for the foraging/raiding process is motivated by the lack of reproducibility in experiments and the inability to isolate individual factors in the ant dynamics that may reveal the true underlying dynamics. Previous mathematical models have implemented many varying approaches to investigate the dynamics of the ant raiding process. Past approaches include continuum PDEs [3, 43, 63, 75, 76], agent-based models through systems of coupled ODEs [7, 11, 24, 47, 57, 74], and lattice models where ants move on a discrete grid [26, 28, 65].

Continuum PDE models forming a system for the ant density and chemical pheromone have shown great qualitative agreement with foraging behavior in ants and allow for rigorous mathematical analysis [2, 3, 54], but the microscopic details of those interactions are lost in the process. Individual-based models have successfully captured individual ant movement by prescribing how each member of the colony behave—even going so far as to separate ants into foragers and returners giving each similar dynamics but different motivation for movement [57]. Furthermore, these models allow the ants themselves to lay the directed pheromone trail to attract others to a given location [11, 57]. These individual-based models also show local traffic dynamics analogous to pedestrians at a crosswalk leading to optimized flow of traffic and resources [24] as seen experimentally [32]. Prior lattice models such as Denuebourg et al. were pioneering for trail formation by using a probabilistic framework to derive multiple trails where ants would pick their next direction based on a biased probability distribution [28]. This was guided by experimental observation and helped explain diverging patterns at two branches of a land bridge where ants communicate only via pheromone [26]. Lattice models have also been used to show that ants highly optimize their trail and dynamics while raiding [65]. Each of these mathematical approaches has revealed interesting features of the raiding process, but none has been dedicated to simultaneous trail formation at multiple food sites.

The primary focus of this work is to develop deeper understanding of ant foraging dynamics by developing a model that incorporates the best features of the lattice-based and agent-based prior approaches. While building upon recent work, the model here is designed to study trail formation in the presence of multiple food sources, which present many interesting questions with biological and mathematical implications. We then rigorously investigate the model through numerical means and reveal underlying properties through the analysis of the corresponding macroscopic PDE formulation. For example, in the presence of a colony’s discovery of multiple food sources, how does a colony allocate its members to retrieve food in an efficient way? One challenge about studying this problem experimentally is that observations suggest that different food sources elicit different recruitment responses indicating that the ants rely on their ability to detect pheromone gradients [40].

We emphasize that our model is intended to be as universal as possible whilst remaining realistic. The primary mechanism of collective behavior in our model is chemotaxis. This is not specific to ants, as it can just as easily be applied to study bacteria [8, 20, 45] and slime molds [13, 23, 72], amongst many other organisms. However, we apply facets of chemotaxis to unveil the simplest possible manner by which emergent spatiotemporal ordering (trail formation) may occur in ants. Our lattice model framework is thus easily amenable to study other organisms of interest as well. In Section 2, we introduce a novel lattice-based model for ant movement in a domain. The primary new feature is the incorporation of principles from agent-based models for ants (e.g., [7,57]) such as direct pheromone deposition by the returning ants at exponentially decreasing quantities as an individual moves away from the food source. In addition, the pheromone concentration is governed by a reaction-diffusion PDE first introduced in [57]. The key assumption is that foraging and returning ants are motivated by different factors and therefore are guided by different rules for their dynamics. Once the lattice model is established, we derive the corresponding macroscopic PDEs governing ant density analogous to past approaches (e.g., [11]). In Section 2.1, we provide details of the computational setup of the numerical simulations for the lattice model and the macroscopic PDE model. In Section 3.1, we perform linear stability analysis on the macroscopic PDE model. We establish a key result that a uniform spread of ants in a domain with a food source is unstable and, furthermore, show numerically that the stable observable state is in fact trail formation. The model is then used to derive enhanced understanding of the multiple food source case which has not been studied extensively in the literature. Understanding this scenario is essential to forming more complete knowledge of ant raiding behavior and provide insight into collective dynamics of social insects as a whole. We conclude in Section 4, where we highlight new biological understanding derived from modeling and simulation predictions. We also setup potential future avenues of investigation.

## 2. Bio-inspired Lattice model

There are challenges related to modeling ant foraging dynamics and trail formation. Namely, the trail formation can depend on model parameters such as distance from the nest, density of the food source and the size of the food source [40]. We forgo the latter of these issues by assuming that each food source contains an infinite reservoir of food. In the following we describe a simple lattice model capturing essential features of ant foraging dynamics whilst forgoing more realistic microscopic details. We find that our simple lattice model produces results akin to what is seen in more complicated agent-based models [57]. Our model is also amenable to analysis at cost of some fidelity to reality.

### 2.1. Lattice Model Description

We model the general terrain as an *M × N* ⊂ ℕ ^2^ lattice and the ants as *n* ∈ ℕ particles hopping along the lattice nodes. We assume that the timescale of trail formation is small enough (approximately 4-8 hours from biological observation of army ants *E. burchellii* [61, 62]) that we ignore births and death in the colony. There is no direct communication between individuals, but rather a response to a chemical pheromone gradient if present represents the only means of (indirect) communication. We designate a single site as the nest, **y**. Initially, all *n* ants occupy that designated site. To understand how the location of food sources relative to the ant nest affects spatiotemporal structure of ant motion, we also randomly designate 𝒩 ∈ ℕ sites as food sources. We assume the colony of ants are self-propelled particles represented by a set of points {**x**_*i*_}, *i* = 1, …, *n*, where **x**_*i*_ ∈ [1, *M* ] *×* [1, *N* ]. Each point can be thought of as the location of the center of mass for an individual ant. We assume the boundaries are reflecting for ants to maintain a fixed population.

Ant motion is subdivided between two types of ants: (i) foraging ants and (ii) carrying ants. Foraging ants are those that are searching for a food source. They undergo a random walk modeled as a Brownian motion [21, 53]. That is, a foraging ant at a given site moves to any adjacent site in its Moore neighborhood with equal probability. In particular, the angular deviation between fixed time steps was measured experimentally in Pharaoh ants and shown to be well approximated by a normally distributed random variable around the nest [10]. At each time step an ant picks a new site to move to based on this probability and thus all ants move with a constant speed. A foraging ant becomes a carrying ant once it locates a food source. Its dynamics are no longer Brownian. Upon reaching the nest, a carrying ant again becomes a foraging ant.

Recent works have incorporated two pheromones into modeling frameworks: one where ants carrying food mark a food source [25, 32] and another where foraging ants mark the location of the nest [41,55,66]. We assume throughout this work that carrying ants know the location of their nest and we choose to forgo the second pheromone in the model. Thus, we only implement the net result of the second chemical deposition by assuming that individual returning ants take the direct path home toward the nest as has been observed experimentally in [15, 49, 50, 77]. Here it was observed experimentally that even obstructing ants by imposing barriers after a food source is located does not dissuade them from following the most direct path home. This suggests the pheromone alone may aid in finding the nest, but a more complex neurological process may contribute as well. In these works, it is noted that ants can follow landmark routes and recognize locations to navigate. The evidence suggests that ants use path integration and their collective knowledge of complex outbound routes to triangulate the terrain and return home directly. Ants do not use complicated path integration akin to humans, but rather use an approximation accounting for navigational errors [49].

To recruit additional ants to the food source a given ant found, a carrying ant secretes pheromone in exponentially decreasing amounts as it travels to its nest to form a chemical trail from the nest to the food source. Though decreasing we assume ants keep secreting pheromone until they reach the nest in contrast to another recent work that has considered ants having a finite chemical supply until they “recharge” at the nest site [47]. The physical response of an ant to a chemical stimulus is well documented [17,24,51]. Following Weber’s Law, foraging ants can then use their antennae to analyze the local concentration of pheromone and preferentially travel against the concentration gradient to reach a food source more efficiently. Recent models assume that to impose Weber’s law the ant’s response rate only depends on the pheromone concentration at the tips of the antennae referred to as *tropotaxis*. The rate of pheromone secretion is assumed to diffuse and decrease exponentially with a carrying ant’s distance from the food location [24,56]. Pheromone deposition and trail laying are modeled by a two-dimensional reaction-diffusion process for the chemical concentration *c*(**x**, *t*) first introduced in [57]:

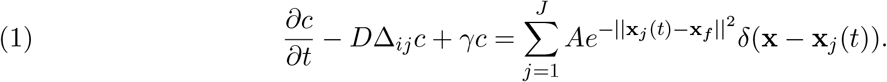

Here **x**_*f*_ is the location of the food, *J* is the number of carrying ants, Δ_*ij*_ is the discrete Laplacian, *D* is the diffusion coefficient controlling the rate at which the pheromone spreads, and *γ* is the evaporation coefficient that ensures an exponential decay of the pheromone in time. The coefficient 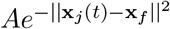 represents the amount of pheromone deposited at time *t* and decays as a carrying ant moves away from the food source. This decrease is needed to ensure that the proper gradient forms due to the competition with diffusion. The initial distribution of chemical is taken as zero so there is no pre-defined directional preference. We prescribe homogeneous Fourier-type boundary conditions for the pheromone so that we have

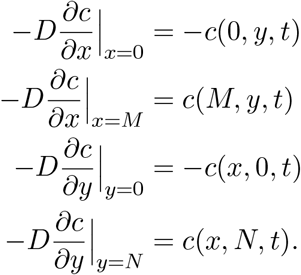

Hence, Fickian flux along the boundary is preserved. That is, along the boundary, the chemical continues to flow with its concentration gradient.

### 2.2. Simulation

On each time step, we select a random site on the lattice. If the site is vacant, we continue randomly selecting a site until an occupied lattice site is selected. One of the ants at that locale is randomly selected to move to one of its neighboring sites according to rules described below. We define *n* such ant movements as one unit of time. Initially, all ants are located at the nest, **y**. At all points in time, we track foraging ants and carrying ants at each location. One can find the simulation parameters and the corresponding biological values taken from experimental observation in Table 1. We describe the simulation in detail in the following.

**Table 1.** Values used in simulation for dimensionless biological parameters.

| Parameter | Non-dim Value (Sim) | Physical Description | Biological Value |
| --- | --- | --- | --- |
| $L$ | 1.0 | Effective ant length/lattice spacing | 1 cm |
| $dt$ | .001 | Time step | 1 sec [3, 57] |
| $N$ | 1000 | Amount of ants in a given colony | $10^2 - 10^6$ (supercolonies [7]) |
| $D$ | 10.0 | Pheromone diffusion coefficient | .01 cm<sup>2</sup>/sec [18, 24] |
| $\gamma$ | .001 | Pheromone degradation coefficient | 1/300 sec<sup>-1</sup> [24, 57] |
| $A$ | 1.0 | Amount of pheromone deposited | $1.1 \times 10^{-4}$ g cm<sup>-2</sup> [18, 24] |
| $\varepsilon$ | $10^{-6}$ | Pheromone conc. detection threshold | Estimated |

#### Foraging Stage

During this stage all ants undergo Brownian motion, so that when an ant is selected to move, it moves into any one of the adjacent sites in its Moore neighborhood with equal probability [3, 11, 63]. This continues until one of the ants finds a food source.

#### Pheromone Secretion

Once an ant finds a food source, **x**_*f*_, it immediately starts secreting pheromone which attracts other ants [42, 66]. The dynamics of the pheromone concentration field, *c*(**x**, *t*), are simulated by Eq. (1). We solve Eq. (1) with a Forward Euler discretization and take the numerical time step Δ*t* = 0.001 units and spatial step Δ*x* = 1 unit.

#### Trail Formation Stage

We assume that ants that reach a food source begin carrying food. Such carrying ants make a beeline (direct path) for the nest [49, 77, 78]. That is, when a carrying ant is selected to move during a simulation, it moves to the first adjacent site encountered when extending the vector **y**−**x**_*f*_ from the center of the site where the carrying ant is selected. The actual movement of a carrying ant will not resemble a beeline on the lattice, but rather an approximation and therefore the lattice model accounts for short length scale randomness in the returning trajectory. However, as lattice size (number of lattice points) is increased, this method will result in a more direct path trajectory for carrying ants returning to the nest.

Foraging ants subject to the pheromone concentration field now undergo a biased Brownian motion. The probability that a foraging ant moves to an adjacent site in its Moore neighborhood is weighted according to the concentration of pheromone at that site relative to the pheromone concentration at the current location of the ant. This is consistent with recent studies showing indirect decision making by individual ants in response to local pheromone gradients (seen both in experiments of [24, 48] and theoretical studies [51]). To implement this, we compute the difference in pheromone concentration at the current location of the ant, **x**_*c*_, and the pheromone concentration at a given adjacent site in the Moore neighborhood of the ant, **x**_*a*_. Here, *a* ∈ *A* and *c* ∉ *A*, where *A* is the set of locations in the Moore neighborhood of the ant selected to move. Thus, **x**_*c*_ ≠ **x**_*a*_.

Let Δ*c*_*a*_ *≡ c*(**x**_*a*_, *t*) *− c*(**x**_*c*_, *t*). We assign a weight, *W*_*a*_, to each site as follows:

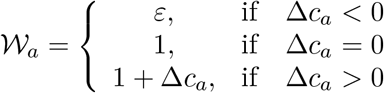

The probability that a foraging ant subject to a pheromone concentration field moves from **x**_*c*_ to **x**_*a*_ is then

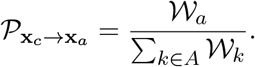

Thus, a foraging ant subject to a pheromone concentration field preferentially moves against the largest pheromone concentration gradient, whilst still having a nonzero probability of moving in neutral directions. We assign a small weight *ε*, with 0 < *ε* ≪ 1, describing the probability of an ant to move down the concentration gradient of a pheromone. The *ε* parameter ensures an ant will not get stuck in a local maximum concentration at a lattice point. In this case, every direction would be assigned *ε* and therefore it would pick a random neighboring lattice point to move to. We note that this situation occurred rarely in simulation where the chemical magnitude was extremely small and had no effective on the global macroscopic behavior of the colony. This is consistent with a recent observation in [80] where they found that there is a minimum concentration threshold below which ants cannot sense and therefore would return to a random walk motion.

This framework for foraging ant motion is advantageous because it incorporates the net effect of Weber’s law by taking into account pheromone concentration all around a given ant to describe its individual response [19, 64]. Weber’s law as it pertains to ants roughly states that if *L* is a pheromone concentration on the left of an ant and *R* is the analog on the right, the individual response is proportional to (*L−R*)(*L*+*R*)^−1^[4]. Weber’s law as applied to ant movement was shown to hold experimentally in a convincing work [52]. Our model captures the net effect of Weber’s law without explicitly encoding it. Instead, Weber’s law emerges from the intrinsic dynamics.

Another advantage of this framework is its flexibility and amenability to alternative modeling frameworks. There have been other mechanisms proposed for how pheromone fields affect ant dynamics. For example, in [47], the authors assume that ants *turn toward* directions of greater pheromone concentration rather than *moving toward* the same direction. Our biased random walk formulation essentially encapsulates both modeling paradigms by incorporating a macroscopic perspective of the underpinning effects of pheromone concentration upon ant motion. The bias towards the highest pheromone concentration can be construed as simply turning toward the local maximum of pheromone whilst still maintaining the ability to detour away from the pheromone source.

*Remark*. We note here that numerical solution of Eq. (1) occurs concurrently with updating ant positions. Because ant position updates occur on a unit time scale and Eq. (1) updates occur on a time scale prescribed by the time step Δ*t*, care must be taken in implementing the simulation. Once Eq. (1) becomes active at the arrival of an ant to a food source, we first updating the PDE and then updating ant positions.

We also note that the results presented for the stochastic model herein correspond to large population size systems. Trail formation in small population stochastic systems can be more challenging since pheromone secretion can fluctuate. However, in the large population size regime, pheromone secretion contracts to the mean, and the formation of trails is better seen and understood.

### 2.3. Lattice Model Results

More realistic models of foraging ant dynamics show that following a transient phase wherein the distribution of ant positions is approximately Gaussian, ants form a robust trail from the nest to a located food source [24, 32, 36, 38, 39, 52, 57, 68]. In the following, we validate our model against the one presented in [57] and then show that our model using parameters in Table 1 can produce more sophisticated results that capture foraging ant behavior seen in experiment or nature. The beauty of our simple model is that it coarse-grains over microscopic underpinnings described in other models to capture the macroscopic picture but nevertheless captures details described in more specific models (see [4, 35, 47, 73], for example). We validate against [57] due to the direct relation between our model and the one presented there. To consider the lattice model (or any model of ant dynamics and trail formation) a success it must at minimum capture three distinct regimes: (i) the colony forages within an area of a given size around a central nest, (ii) the model has the ability to recruit ants to any food sources discovered, and (iii) the model exhibits complex macroscale patterns of trails emanating from the nest [55, 80].

#### Validation

To validate our model, we compare our results with those presented in a more realistic agent-based model [7, 57] and experiment (e.g., [24]). In [57], Ryan presents a detailed agent-based model describing ant foraging dynamics. There, before any food is found, the distribution of ants about the nest is approximately Gaussian. Once a food source is found, pheromone secretion ensues and there is a rapid transition to a state where an ant trail connecting the nest to the food location forms. Ryan also showed that the interior of well-formed trails consisted of returning ants and the exterior of foraging ants.

In Figure 2, we show sample simulations of our lattice model that depict the same behavior. Initially, the distribution of foraging ants about the ant nest is approximately Gaussian, stemming from their dynamics being a discrete Brownian motion (Figure 2a). Once a food source is located and pheromone secretion is underway, ants begin to self-organize and form a trail connecting the nest to the food source (Figure 2b). Eventually, the vast majority of the ants partake in a well-formed trail between the nest and the food source (Figure 2c).

**Figure 1.**
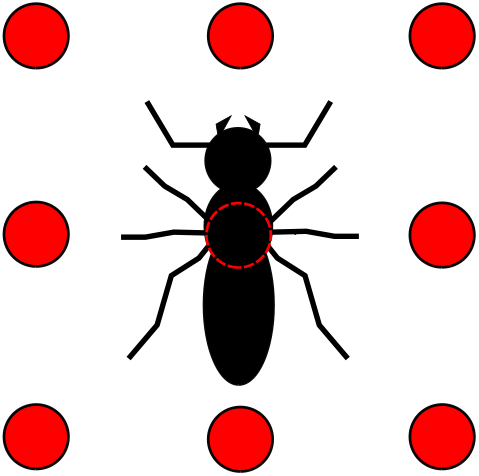
Illustration of an ant on the lattice and the positions of its neighboring sites in its Moore neighborhood. We omit the details of an individual ant’s body such as its three components and individual appendages/antennae in the model. Rather we simulate the change in a given ants center of mass across the lattice network. This remains faithful to the biology by considering many ants in a large computational domain where the individual microscopic details becomes less relevant to understanding the macroscopic behavior.

**Figure 2.**
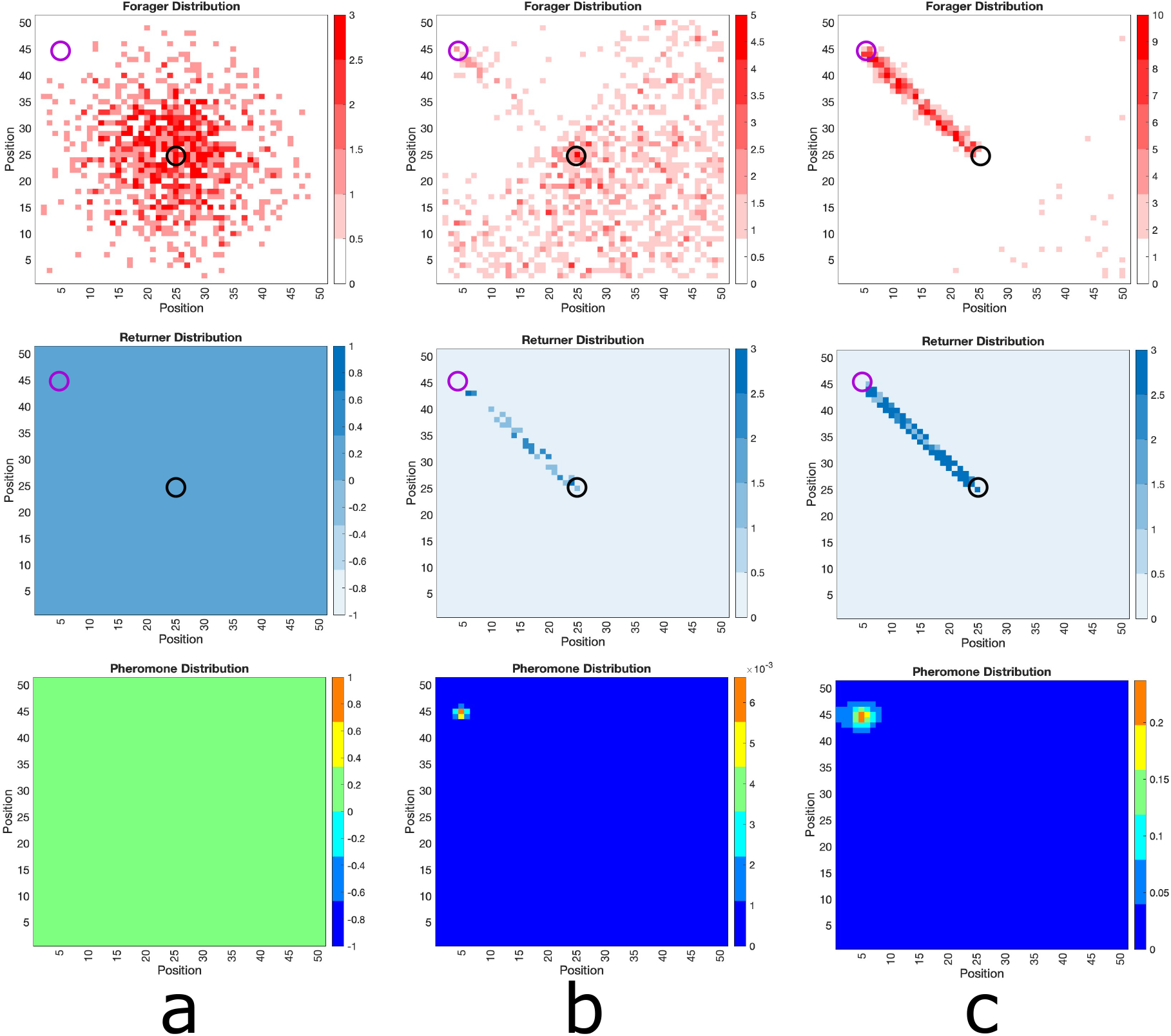
Foraging Ant Dynamics: One Food Source. Top row: foraging ant distribution. Middle row: returning any distribution. Bottom row: pheromone distribution. **(a)** The foraging ants distribution is approximately Gaussian around the nest (center black circle) until food source (purple circle) is discovered, whence pheromone is released as returners move toward the nest. **(b)** The pheromone excretion initiates emergent self-organization of the ants into a trail connecting the nest to the food source. **(c)** The trail is well-developed. Due to the fact that the food at the source does not diminish, this trail continues forever.

Furthermore, returning ants in a well-formed trail follow a direct path (Figure 2c, middle row), whereas foraging ant locations (top row) are distributed around the main trail. Hence, returning ants are definitively found in the interior of well-formed trails whereas foraging ants can be found on the exterior. Thus, our simple lattice model reproduces the results of the more realistic agent-based model of Ryan. We also note here that the results are consistent with experimental results [24, 52]. In particular, the foraging trail is wider resulting in more ants on the outside of the central trail lane. This exactly matches the qualitative observations of ants in the experimental observations of Couzin et al. [24] and it is remarkably captured in our simple lattice model Fig. 2c.

#### Multiple Food Sources

The detailed agent-based model presented in [57] was lacking in one regard: when multiple food sources were present, the model did not allow for the persistence of multiple ant trails connecting the nest to each food source. Experimental studies, however, clearly show that ant colonies establish multiple ant trails in the presence of multiple distinct food sources [16].

Our simple lattice model allows for the persistence of multiple ant trails in multiple food source environments. In Figure 3, we show snapshots of foraging ant dynamics in an environment consisting of two equidistant food sources from the ant nest. Initially, there are no returning ants and the foraging ant distribution is approximately Gaussian (Figure 3a). Once the food sources are found, the ants begin to self-organize into two distinct ant trials (Figure 3b) and then establish well-formed trails connecting the nest to the food sources (Figure 3c).

**Figure 3.**
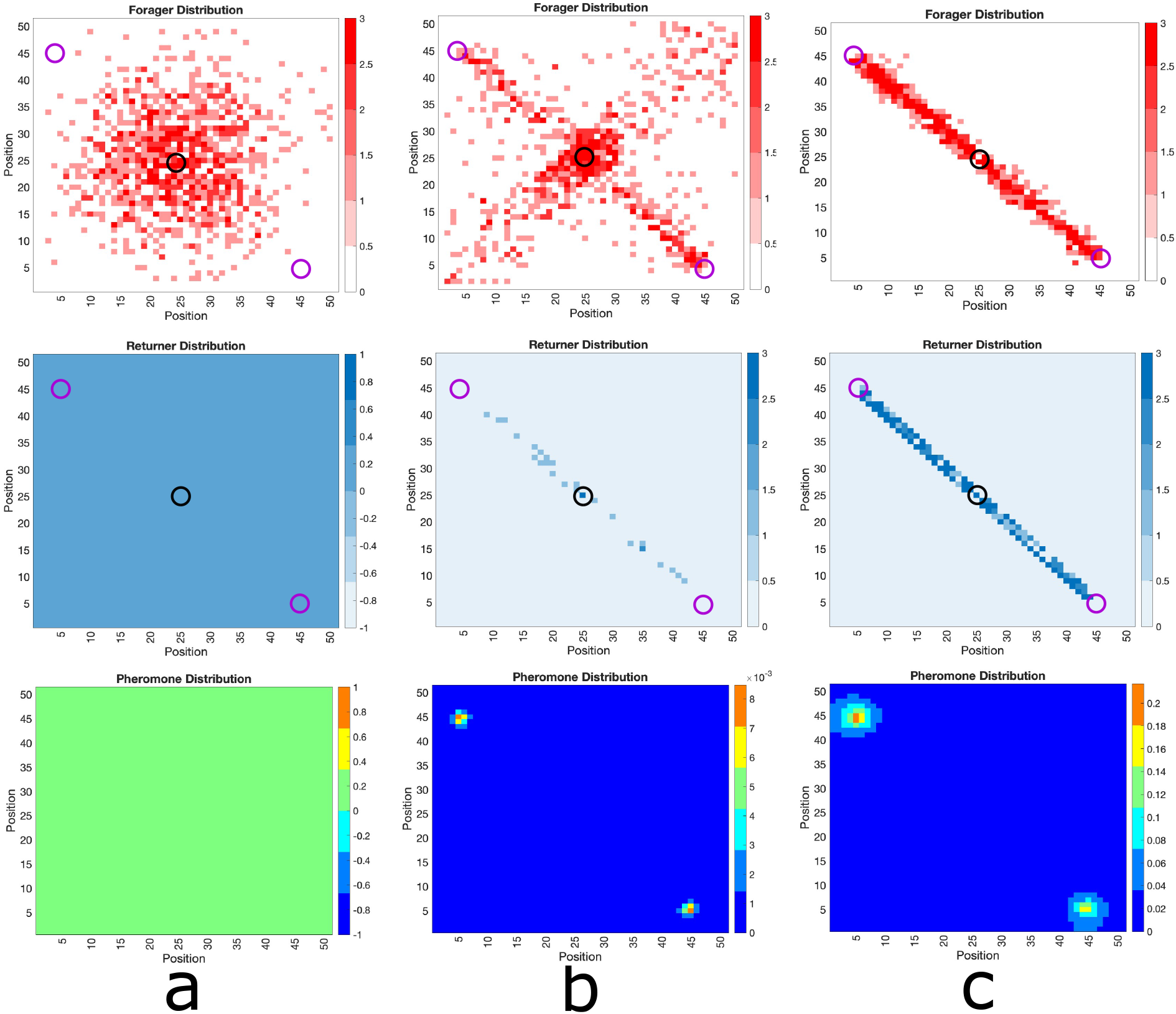
Foraging Ant Dynamics: Two Equidistant Food Sources. Top row: foraging ant distribution. Middle row: returning any distribution. Bottom row: pheromone distribution. **(a)** The foraging ants distribution is approximately Gaussian around the nest (center black circle) until the food sources (purple circles) are discovered, whence pheromone is released as returners move toward the nest. **(b)** The pheromone excretion initiates emergent self-organization of the ants into trails connecting the nest to the food sources. **(c)** The trails are well-developed. The food source does not diminish and there is no asymmetry in the properties of the food sources relative to the ant nest, both trails continue forever.

If two food sources are present but not equidistant from the ant nest, the results are similar (see Figure 4). The main difference is that the state in which multiple trails connecting the nest to the distinct food sources manifests as a *quasistationary* state (Figure 4c) before all ants converge to the trail connecting the nest to the nearer food source (Figure 4d). In this context, the disparity in distances to distinct food sources renders the use of ants to obtain food from farther away inefficient. In this situation, the model predicts that ants will preferentially collect all the food from the nearer food source before traveling farther for food. Even in the case presented in Figure 4, wherein the farther food source is just one unit farther than the closer food source, the asymmetry in the system is sufficient for the ant colony to focus all of its population on the closer food source. This can be explained simply by appealing to diffusion correlation length of the pheromone signal. The closer food source manifested a stronger pheromone signal near the nest, which in turn resulted in stronger ant recruitment. The recruitment then facilitated a stronger pheromone gradient signal connecting the nest to the closer food source.

**Figure 4.**
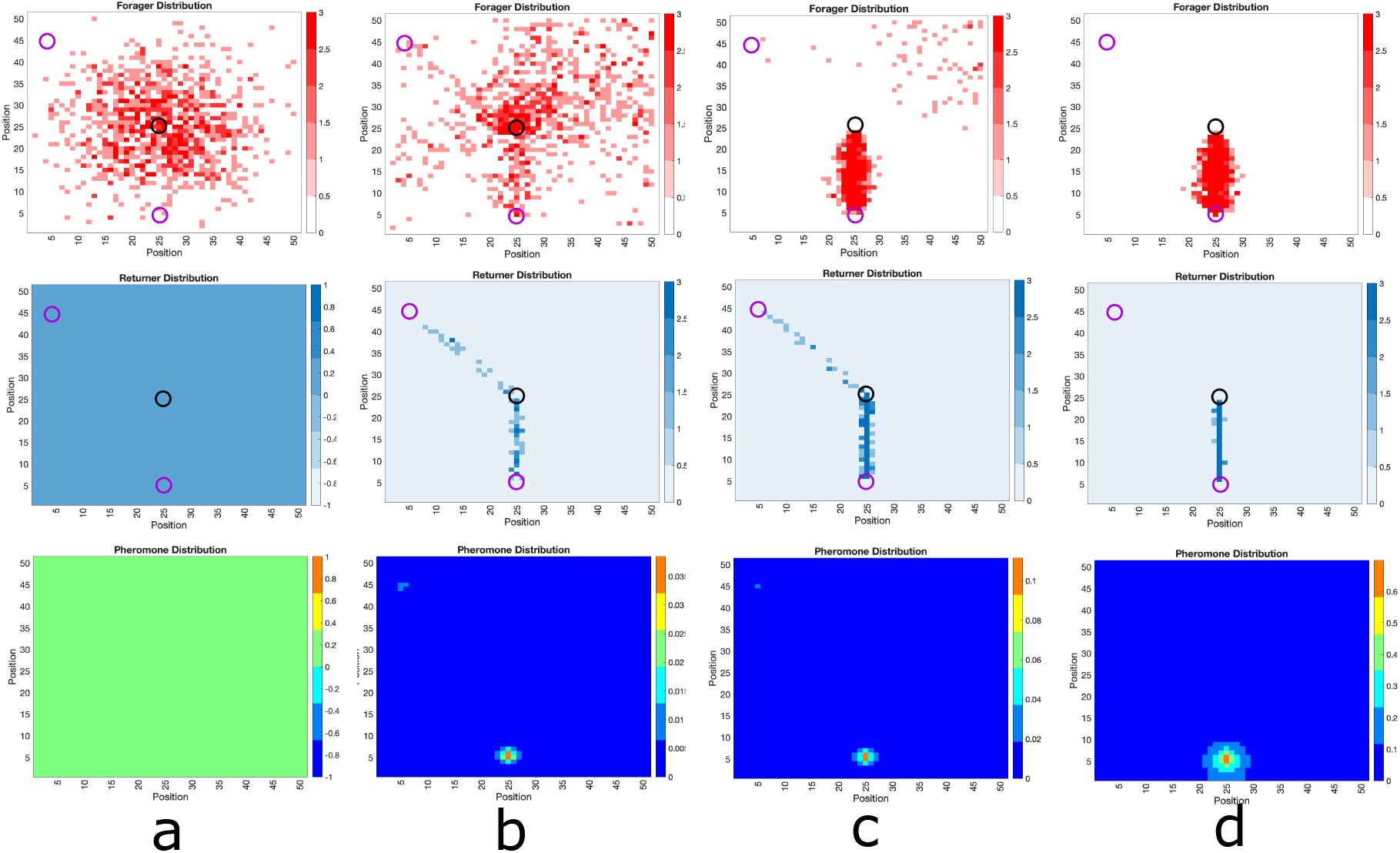
Foraging Ant Dynamics: Two Food Sources: One closer, one far. Top row: foraging ant distribution. Middle row: returning any distribution. Bottom row: pheromone distribution. **(a)** The foraging ants distribution is approximately Gaussian around the nest (center black circle) until the food sources (purple circles) are discovered, whence pheromone is released as returners move toward the nest. **(b)** The pheromone excretion initiates emergent self-organization of the ants into trails connecting the nest to the food sources. **(c), (d)** Both trails are well-developed at short time scales, but because it is less efficient to send ants to the farther food source and less ants visiting leads to the pheromones on that trail dissipating faster. Therefore, eventually all ants converge to the trail connecting the nest to the closer food source. Because food at the source does not diminish, this trail continues forever.

This latter effect is observed in “winner-take-all” processes in biology, wherein identical targets compete for a limited resource of nutrients. The emergence of some asymmetry in the system eventually pushes all the resources to be held by one of the targets. One example of such a process is neurite polarization, wherein one of several identical protrusions (neurites) emanating from a nascent neuron compete for a common resource (such as tubulin). Due to intracellular and thermodynamic noise, one protrusion collects more resource than others by chance. This single neurite becomes the axon of the mature neuron, while the others become dendrites [6, 59, 70].

Such processes are also observed in natural and synthetic gene circuits that involve mutual inhibition positive feedback loops [1, 9, 58]. A motif of such systems is bistability—wherein at equilibrium one protein persists and the other goes extinct. The persistence of either protein in the system is equally likely, but initial conditions drive the system one way or the other. Such systems are called “toggles” [71], as varying key biophysical parameters can shift the equilibrium to a state where the other protein persists. Winner-take-all mechanisms are also present in learning in neural networks [44, 46] and synthetic gene circuits [67], among many other systems.

In this perspective, ants can be construed as resources and the food sources as targets. When there is nothing differentiating the identical food sources (as in Figure 3), ants form longstanding trails to both food sources. When asymmetry in the distances to the respective food sources is present, however, the closer food source eventually collects all the ants.

Thus, our simple lattice model consisting of only essential basic features of foraging ant dynamics produces results that are observed in more realistic agent-based models and in experimental studies. One advantage of the lattice model is that it is simple to implement. Another advantage is that it is amenable to analysis, which we discuss in the next section.

## 3 Macroscopic Probabilistic Model

The lattice model is simple enough so that it is amenable to analysis. To understand the dynamics of the stochastic lattice model we develop a macroscopic equation describing the evolution of probability densities of foraging and carrying ants. Although the simulated system is spatially discrete, the corresponding discrete master equation is cumbersome and unintuitive. In the limit of a very large lattice size and small grid size, the master equation is well-approximated by the corresponding macroscopic PDE. We present the model for a single food source, but it is easily generalized to multiple food sources. Let *p*(**x**, *t*) denote the probability density for the position of a foraging ant at time *t* and let *q*(**x**, *t*) denote the probability density for the position of a carrying ant at time *t*. Finally, let *u*(**x**, *t*) be the continuum analog of *c*(**x**, *t*). Then, the dynamics *p, q*, and *u* can be characterized by

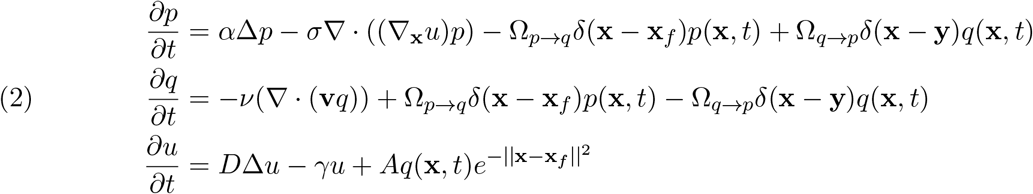

Here, *α* is the diffusion coefficient for the foraging ants and *σ* describes the sensitivity of the foraging ants to the pheromone. Mathematically, *u*(**x**, *t*) can be viewed as a potential. The rates Ω_*p→q*_ and Ω_*q→p*_ are the transition rates of foraging ants becoming carrying ants and vice versa, respectively. The multiplying Dirac delta functions impose that the transitions can only occur at a food source or the nest. The coefficient *ν* describes the velocity of a carrying ant, and 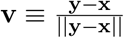 is a unit vector directed at the nest. The dynamics for *u*(**x**, *t*) are continuum analogs of the dynamics for *c*(**x**, *t*) shown in Eq. (1). We prescribe homogeneous Neumann conditions for *p*(**x**, *t*) and *q*(**x**, *t*) so that, like the lattice simulation, ants are conserved. For *u*(**x**, *t*) we prescribe homogeneous Fourier-type conditions. For initial data, we prescribe *p*(**x**, *t*) = *δ*(**x** − **y**) and zero for *q*(**x**, *t*) and *u*(**x**, *t*).

The solutions to Eqs. (2) will describe the probability densities for a single foraging ant (*p*(**x**, *t*)) and a single carrying ant (*q*(**x**, *t*)) and will also describe the densities of foraging and carrying ants in the macroscopic limit (i.e., the large number of ants limit).

### 3.1. Macroscopic PDE Model Results

Here we first establish that the results of the macroscopic model coincide with the lattice model results before performing a linear stability analysis showing that the presence of food sources is the key factor in establishing ant trails.

#### Comparison with Lattice Model

In Figures 5–7, we show solutions of Eq. (2) for three distinct situations corresponding to the results shown in Figures 2–4. The macroscopic PDE model solutions coincide with lattice model simulations in all cases. In Figure 5, we show foraging ant dynamics in an environment with a single food source. Here, following a transient Gaussian-distribution phase (Figure 5a-c), all ants converge to a single trail connecting ant nest to the food source. In Figure 6, we show foraging ant dynamics in an environment with two food sources equidistant from the ant nest. We see here that the stationary state consists of two trails that form connecting the nest to the food sources. Finally, in Figure 7, we show foraging ant dynamics in an environment with two food sources, each located a different distance away from the nest. As in the lattice simulations (see Figure 4), following a transient quasistationary phase where two trails form, the stationary state consists of a single trail connecting the nest to the proximal food source.

**Figure 5.**
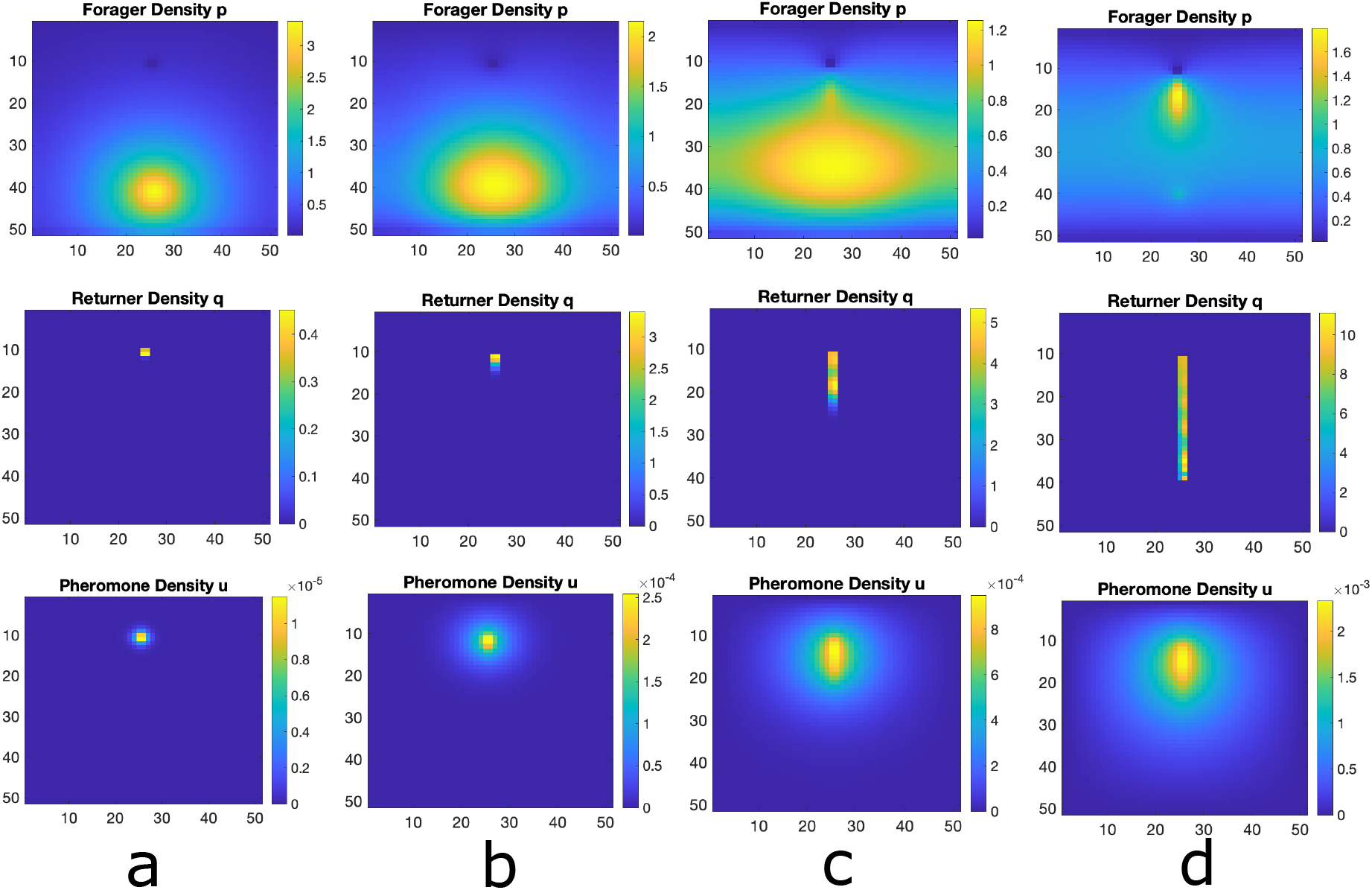
Numerical Simulation of Macroscopic PDE System. Simulated Eqns. (2) for forager density *p*, returner density, *q*, and pheromone density, *u*, using the Alternating Direction Implicit Method (ADI). a) Initial configuration with a bump function of foragers and essentially no returners or pheromone, b) food source is discovered and pheromone is released as returners move toward the nest. c) the trail of foragers and returner develops at intermediate times. d) the trail is well developed. This shows that the perturbation from the homogeneous solution discovered analytically above actually leads to the trail formation as at least a quasi-stable state.

**Figure 6.**
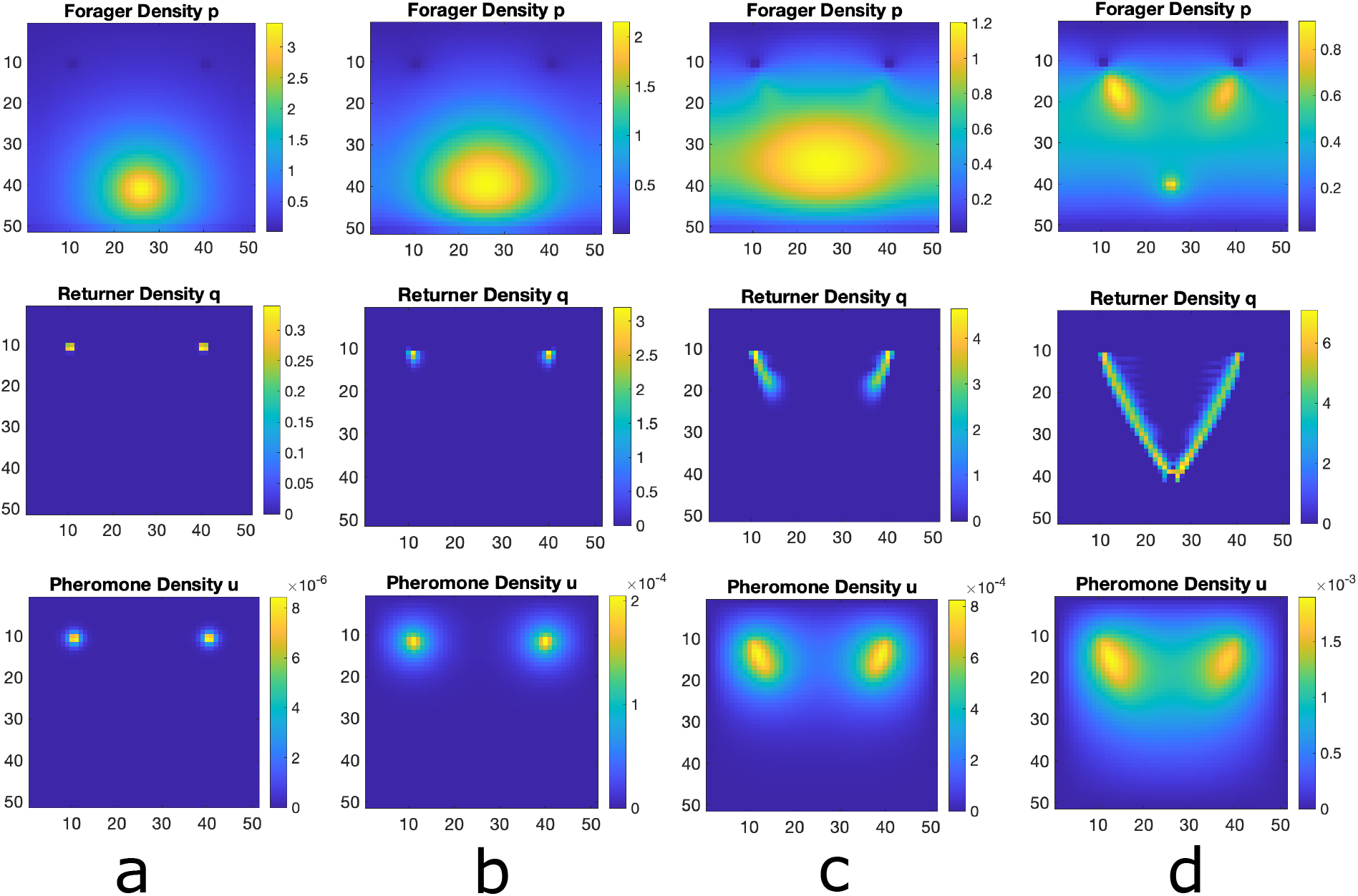
Numerical Simulation of Macroscopic PDE System for Multiple Food Sources. Simulated Eqns. (2) for forager density *p*, returner density, *q*, and pheromone density, *u*, using the Alternating Direction Implicit Method (ADI). a) Initial configuration with a bump function of foragers and essentially no returners or pheromone, b) food source is discovered and pheromone is released as returners move toward the nest. c) the trail of foragers an returner develops at intermediate times. d) the trail is well developed. This shows that the master equation is capable of handling multiple food sources.

**Figure 7.**
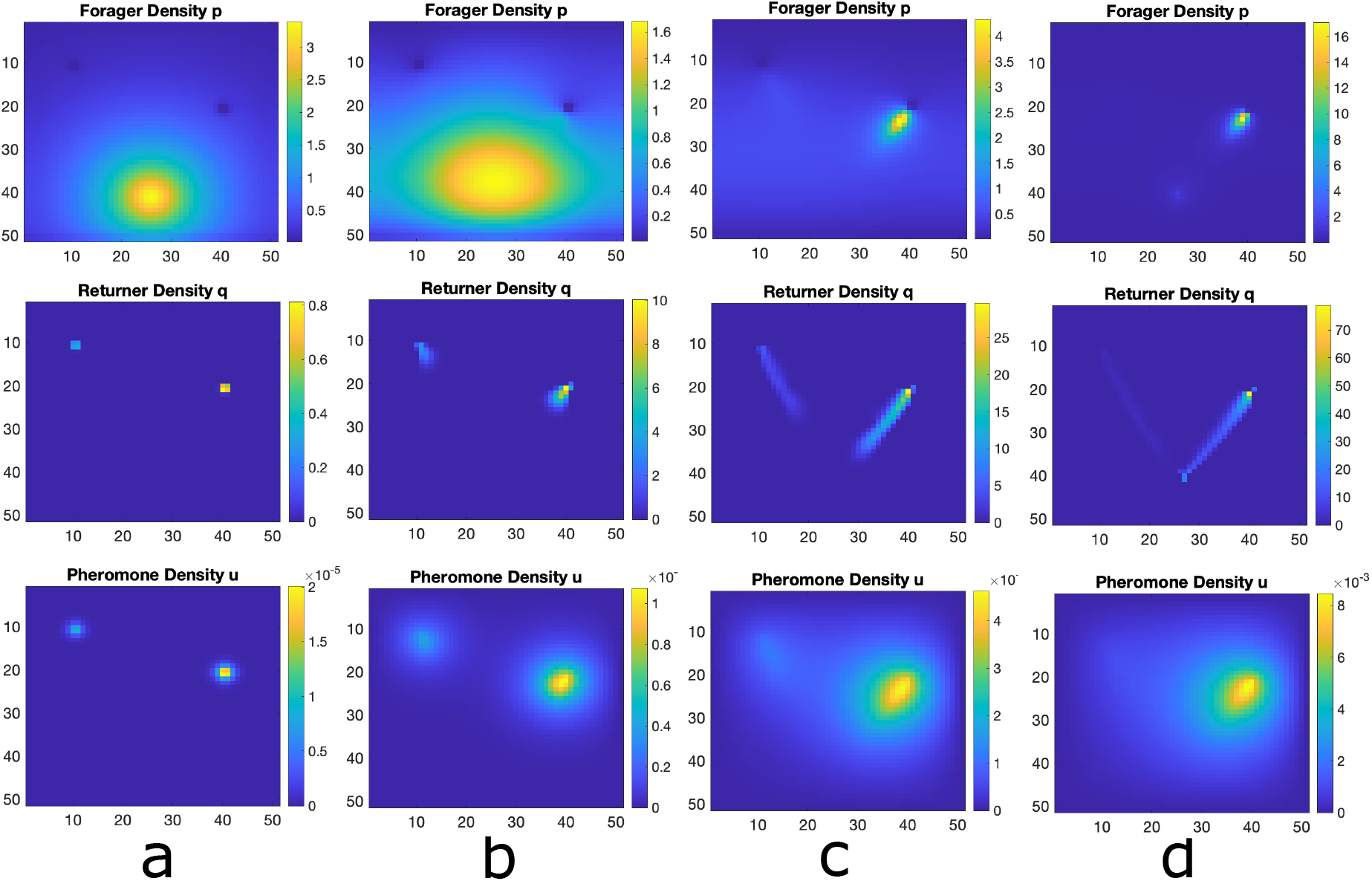
Numerical Simulation of Macroscopic PDE System for Multiple Food Sources at Unequal Distances. Simulated Eqns. (2) for forager density *p*, returner density, *q*, and pheromone density, *u*, using the Alternating Direction Implicit Method (ADI). a) Initial configuration with a bump function of foragers and essentially no returners or pheromone, b) first food source is discovered and pheromone is released as returners move toward the nest. c) the trail of foragers an returner develops at intermediate times to both food sources. d) the trail is well developed at the closer food source and the trail to the farther food source evaporates naturally due to no ant visits.

#### Linear Stability Analysis

Here we show analytically that the presence of a food source in the domain of interest promotes the formation of ant trails. In the absence of food, Eqs. (2) reduce to

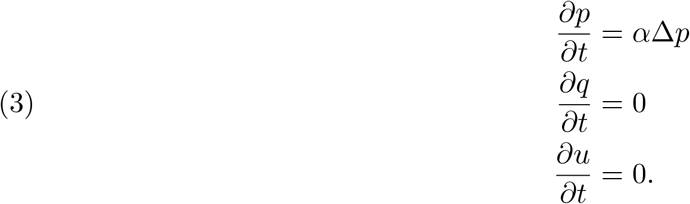

In the absence of a food source, foraging ants never become carrying ants and thus never produce pheromone. Thus, carrying ant dynamics and pheromone dynamics are trivial. Initial data then yield the homogeneous equilibrium 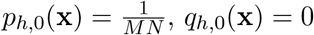 and *u*_*h*,0_(**x**) = 0.

We next show that the presence of food promotes a nonhomogeneous equilibrium to emerge. We equate this emergence with the formation of an ant trail from the nest to the food source. To do this, we invoke a mean-field approximation upon Eqs. (2) so that coefficients are space-independent.

This yields the equations

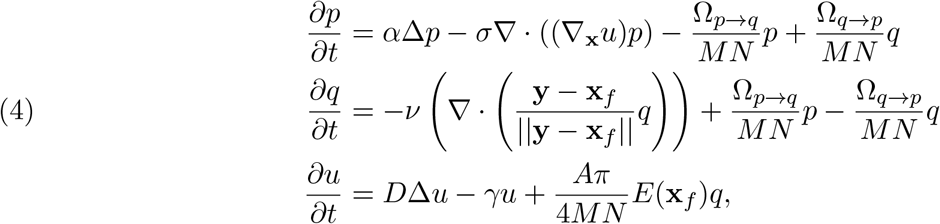

where we take **x**_*f*_ = ⟨*x*_*f*1_, *x*_*f*2_⟩ and *E*(**x**_*f*_) *≡* (erf(*x*_*f*1_) − erf(*x*_*f*1_ *− N*))(erf(*x*_*f*2_) − erf(*x*_*f*2_ *− M*)). A homogeneous equilibrium for Eqs. (4) is given by

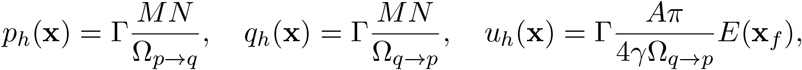

where 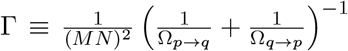 Observe that by taking the limits Ω_*p→q*_ *→* 0, Ω_*q→p*_ *→ ∞*, and *E*(**x**_*f*_) *→* 0, we obtain the homogeneous equilibrium of the system without food: 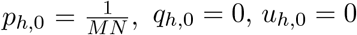 The equations are thus self-consistent.

We next show that the presence of a food source promotes a nonhomogeneous equilibrium to emerge. We equate this emergence with the formation of an ant trail from the nest to the food source. Following a common approach, we will linearize Eqs. (4) about the homogeneous equilibrium and study the spectrum of the resulting linear operator. We consider the case where *M, N* are large relative to the distance between the nest and food source so that boundary effects are negligible for trail formation dynamics. This is a reasonable assumption because the trail formation in the mechanism studied in this paper is fueled through diffusive coupling. Large distances render diffusive coupling inefficient. Hence, we consider Eqs. (4) in the unbounded domain ℝ ^2^.

We assume a solution of the form

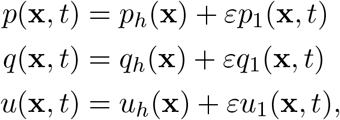

where 0 *< ε ≪* 1 is a small parameter quantitating the perturbation imparted on the homogeneous solution. We look for plane wave solutions with wavevector 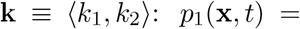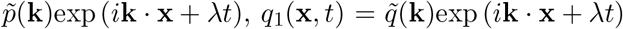 and *u*_1_(**x**, *t*) = *ũ*(**k**)exp (*i***k. x** + *λt*). Substituting into Eqs. (4) and collecting *O*(*ε*) terms yields the linear system **A**|*p*⟩ = |0⟩, where

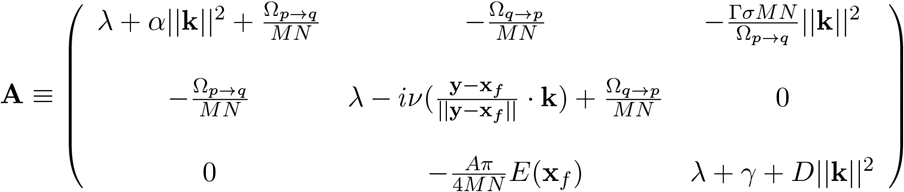

and 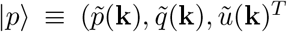. To ensure |*p*⟩ is nontrivial, we set det(**A**) = 0. This yields an eigenvalue problem yielding a dispersion relation between *λ* and **k**. The goal is to determine conditions under which Re[*λ*(**k**)] *>* 0, meaning the homogeneous solution is unstable under a perturbation with the specified wave vector.

Although in theory *λ*(**k**) is computable, the expression is long and unwieldy. We use MATLAB to compute the dispersion relation. The dispersion relation is symmetric with respect to *k*_1_, *k*_2_. Thus, in Figure 8, we show the projection of the dispersion surface onto *λk*_1_–space. Under the prescribed parameters, clearly *λ*(**k**) *>* 0 for a range of wavevectors, indicating instability of the homogeneous solution to low frequency perturbations. The homogeneous solution is stable under high frequency perturbations. However, the instability to low frequency perturbations is what we interpret as the onset of ant trails (nonhomogeneous solution).

**Figure 8.**
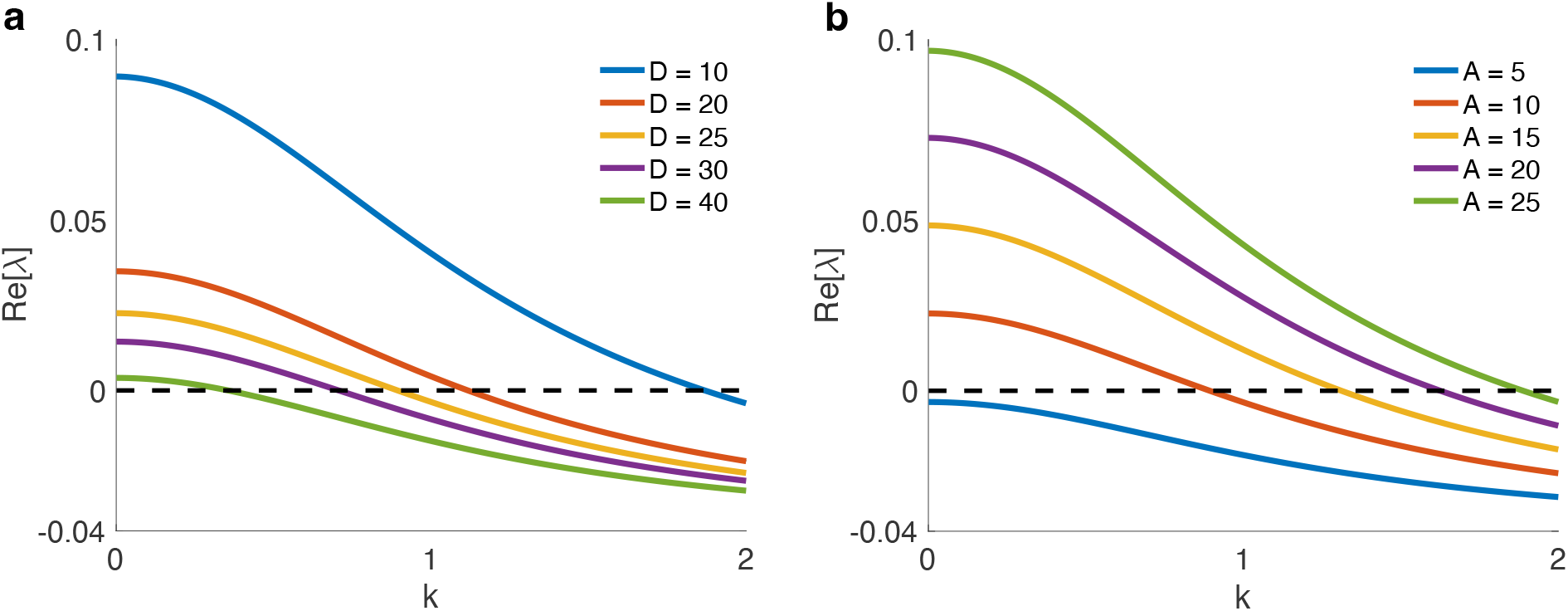
How *λ* varies with wavevector **k**. This is a projection of the dispersion relation onto one of the components of **k**. The dispersion relation is symmetric with respect to the wavevector components. **(a)** Increasing the diffusion rate of the pheromone signal stabilizes the homogeneous steady state. **(b)** Increasing the quantity of pheromone secreted per unit time destabilizes the homogeneous steady state.

To investigate this further, we show in Figure 8a how the dispersion relation changes when *D*, the diffusion coefficient of the pheromone signal, is altered. We find that as *D* is increased, the homogeneous steady state is stabilized. This is easy to understand: when *D* is large, diffusion dominates pheromone dynamics and diminishes any observable gradient in the pheromone concentration field. With a uniform pheromone distribution, ants will be unable to form trails.

In Figure 8b, we show how the dispersion relation changes as *A*, the quantity of pheromone secreted by an ant per unit time, is altered. We find that as *A* is increased, the homogeneous steady state is destabilized. If not enough pheromone is secreted, then clearly there will not be enough pheromone for ants to sense and modify their motion. Indeed, for *A* = 5, we observe an unconditionally stable stationary state. Trails form only when enough pheromone is secreted into the environment.

Thus, our macroscopic PDE model results coincide with the stochastic lattice model results. Moreover, the PDE model is amenable to analysis, and it makes explicit the crucial role the presence of food in an environment plays on the onset of emergent spatiotemporal ordering in ants. Linear stability analysis allows us to investigate the impact of various parameters upon the stability of the equilibrium with trails connecting the ant nest to food sources.

## Discussion

We have introduced a novel lattice model describing foraging ant dynamics that captures only basic essential features of forager ant dynamics and forgoes more detailed interactions. Indeed, the driving mechanism we incorporate in our model is chemotaxis and is not specific to ants. But we have shown that it is sufficient to describe complex spatiotemporal ordering in foraging ants. Our model reproduces results produced by more detailed agent-based models, and even extends some results. Specifically, our model admits multiple trails from the ant nest to multiple distinct food sources, improving on previous models which could not sustain multiple ant trails in the presence of multiple food sources.

Interestingly, our model shows strong prejudice when asymmetry is present in the system. Namely, when multiple food sources are present, but are not equidistant from the nest, at equilibrium all ants are in the trail connecting the nest to the nearest food source. The implication is that sending ants to extract food from farther food sources is inefficient. This maps to experimental observations describing the efficiency of ants [77, 78]. Initially, the ants form trails to all the available food sources. This stems from the fact that in the absence of a pheromone concentration field, foragers perform a 2D random walk. The multiple food trails manifest as a quasi-stationary state. Once pheromone from the nearest food source becomes perceptible, all ants converge to the trail forming between the nest and the nearest food source. When food sources are equidistant, ant trails to each food source manifest in the equilibrium structure.

The simplicity of our model allows for analysis. We derived a macroscopic PDE model that makes explicit the crucial role the presence of food sources plays in the onset of ant trails. Furthermore, linear stability analysis of the PDE model shows that the diffusion rate of the secreted pheromone and pheromone secretion rate are key drivers of emergent spatiotemporal ordering in foraging ant dynamics. Thus, the amenability of our lattice model allows us to unveil key biophysical parameters that govern ordering in foraging ant dynamics.

The directness of our model facilitates a number of future extensions we hope to explore. First, we hope to explore the impact of terrain upon trail formation. To incorporate this, we can prescribe space-dependent hopping rates in our lattice model—smaller hopping rates capture more difficult terrain and larger hopping rates represent flat terrain. Second, we hope to explore the impact of competition between distinct ant colonies on trail formation. Third, we hope to investigate the impact of environmental factors (like rain, which can disrupt pheromone gradient formation) and predators upon trail formation. In the latter case, groups of ants are more prone to predator attacks than isolated ants, causing a cost-benefit problem between ants grouping together to acquire food and isolating away from one another to minimize death due to predation.

Finally, we note that other recent work found that ants actively avoid crowded food sources to facilitate trail formation to other food sources [79]. The driving mechanism there was a combination of chemical interactions and individual interactions. We can incorporate individual interactions by modeling the stochastic lattice model as a modification of a TASEP [14, 22, 29, 69] or by including a Lennard-Jones potential [34, 37, 57] into the macroscopic PDE formulation.

## 5. Statements and Declarations

We declare that we have no competing interests.

## 6. Data Availability

All code used to produce figures in this manuscript will be made available to anyone upon reasonable request.

## Acknowledgements

BRK would like the thank his wife, Hajra Habib, and his son, Surya Karamched, for the being the light of his life.

